# Phylogeography of the desert scorpion *Mesobuthus mongolicus* illuminates a route out of Central Asia and Junggar Basin as a corridor for faunal exchange between Central and East Asia

**DOI:** 10.1101/2021.10.31.466646

**Authors:** Cheng-Min Shi, Xue-Shu Zhang, Lin Liu, Ya-Jie Ji, De-Xing Zhang

## Abstract

A comprehensive understanding of the ecological, evolutionary and genetic consequences of climate changes requires integration of information from different geographic regions and ecosystems. However, a clear knowledge gap exists in arid biota of Asian drylands, where climate change and biological evolution demonstrate high regional specificity. We report here our phylogeographic study on the desert scorpion *Mesobuthus mongolicus*, attempting to narrow the gap. Range wide samples for one mitochondrial gene and three nuclear loci were subjected to phylogenetic inferences, molecular dating, Bayesian phylogeographic and demographic reconstructions, and population genetic analyses. Ecological niche modeling was employed to generate models of potential distribution range at present and during the last glacial maximum (LGM). Our results revealed that *M. mongolicus* represents a single coherent lineage that diverged with its most closely related lineage from Central Asia about 1.36 Ma and underwent radiation ever since. Bayesian phylogeographic reconstruction suggested the ancestral population of *M. mongolicus* dispersed gradually eastward through the Junngar Basin to the Gobi region during the late Pleistocene, strengthening the view that the Junggar Basin has constituted an important corridor for faunal exchange between Central Asian and East Asia. We found that the desert scorpion underwent demographic expansion while experiencing range contraction during the LGM. Development of sandy deserts in northwest China incurred by Eurasian aridification might have opened up substantial empty niches that sustained population expansion during the LGM. Our results suggested that organisms’ ecological adaptations have acted as an important determinant for their phylogeographic and demographic responses to past climate changes.

---

Geographic distribution of a species is an outcome of interplay between environmental challenges and its biological adaptation. Given a certain environmental change, it can either pose a threat or provide an opportunity depending on species’ demographic, physiological and behavior adaptational profiles (Garcia et al. 2014). Thus, species might respond differently to climatic changes in an individualistic way (Ackerly et al. 2010; Anadón et al. 2015; Hewitt 2000; Willis et al. 2004). Besides, historical climate changes were expressed variably across geographic regions (Hewitt, 2000), and their impacts were unfolded in multiple dimensions across geographic locations (Garcia et al. 2014). These features make comprehension of the biological and ecological impacts of climate changes a very complex issue that requires to integrate information from a diverse of organisms, ecosystems, geographic locations and climate regimes.

However, studies on such topics are focused predominantly on Europe and North America and essentially on the impact of glaciations (Keppel et al. 2012). In those regions, it is generally well acknowledged that the glacial cycles have played a major role in shaping geographic distributions and reshuffling genetic variations of plants and animals (Hewitt 2000). On the contrary, the arid regions and the role of aridification, a more widespread coeval of the Quaternary glaciation, remain to be fully addressed (Byrne et al. 2008, 2018; Shi et al. 2013).

Drylands, including deserts and semi-deserts, cover about 40% of the Earth’s terrestrial surface, stretching particularly widely in mid latitude Eurasia (Middleton and Sternberg 2013). Those regions encompass a wide variety of physical terrains. However, what all drylands have in common is aridity, expressed as characteristically scarce precipitation and extremely high evaporation. Such climatic conditions have shaped some of most sensitive and fragile desert ecosystems on the Earth (Austin 2011). Since 22 million years ago, aridification has been a surrogate of climatic changes and profoundly shaped the evolution of biota in the Asian interior (Bougeois et al. 2018; Guo et al. 2002). It has been suggested that the phylogeographical effects of aridification in the subtropic hot deserts mirror or contrast, depending on the ecological adaptation of studied species, the glaciation in the Northern Hemisphere (Fujita et al. 2010). However, this conjecture remains to be tested in the cold deserts in the temperate regions, particularly the drylands in Asian interior.

An expansion-contraction (EC) model has been proposed to describe the phylogeographic dynamics of biota in response to the Pleistocene glacial oscillations (Provan and Bennet 2008). EC model has stood extensively empirical corroboration with temperate biota in Europe and North America. For temperate species, the advance and retreat of ice sheets imposed a consequent parallel contraction-expansion of species’ range and population size (Hewitt 2004; Provan and Bennett 2008; Salvi et al. 2014; Schmitt 2007). On the contrary, for cold-adapted taxa, an inverse model has been proposed (Galbreath et al. 2009; Willis et al. 2004). Evidence is accumulating that EC model is also applicable to subtropical arid ecosystems although the underlying driving climatic factor might differ. In those hot deserts, precipitation rather than temperature, is the limiting resource (Noy-Meir 1973), and has exerted more impact on phylogeographic history (Cosacov et al. 2013). For example, Australia’s arid-adapted Grey Butcherbird (*Cracticus torquatus*) experienced population expansions during Pleistocene glacial maxima whereas population of its mesic-adapted relatives remained stable (Kearns et al. 2014).

Here we test the EC model in the Asian cold deserts including the Gobi where both temperature and aridity pose great challenges to biota, using the desert scorpion *Mesobuthus mongolicus* (Birula, 1911) as the studying organism. This scorpion was formerly known as *M. eupeus mongolicus* and has recently been upgraded to an independent species (Kovařík 2019). It represents the easternmost member of the *M. eupeus* complex (Fet et al. 2018; Fet et al. 2000; Shi et al. 2015). Previously, we have shown that the ancestral distribution area of the *M. eupeus* complex is most likely in Anatolia (Shi et al. 2013). Gobi is the largest desert of Asia, covering huge parts of northern China and of southern Mongolia (ca. 1.3 million km^2^). It is located far north than other temperate deserts on a plateau more than 900 m above sea level. As a consequence, the temperature in Gobi can be extremely hot (>50 °C) in summer and extremely cold (−49 °C) in winter (Pfeiffer et al. 2003). Locked by rain shadows created by the lofty Qinghai-Tibet Plateau and the Tianshan and Altai mountains, Gobi receives very scarce rainfall but endures extremely heavy evaporation. On such an ‘edge of life’, however, *M. mongolicus* (Mmon hereafter) thrives (Shi et al. 2007, 2015), providing an ideal model to bridge the knowledge gap in dryland phylogeography. The Gobi populations of Mmon has been proposed as a product of out-of-Central Asia move of its ancestors during the late Pleistocene (Shi et al. 2013). However, the exact route through which Mmon migrated to the Gobi remains to be defined. In the present study, we have tested the EC model by combining genetic analysis and ecological niche modelling of Mmon, with the aim to explore (1) the possible route through which Mmon dispersed out of Central Asia, (2) its demographic pattern casted by past climate changes, and (3) the climatic context that has shaped its phylogeography.

## Materials and Methods

### Sample collection and genetic data

The natural distributional range of Mmon has been determined though systematic field survey with the aid of species distribution modeling (Shi et al. 2015). We sampled 338 individuals of Mmon from 51 sites, including 41 sites in China (C_1_-C_41_, *n* = 274), 8 sites in Mongolia (M_1_-M_8_, *n* = 62), one site each in Tajikistan (TJ1, *n* = 1) and Kazakhstan (KZ1, *n* = 1) (Figure 1). The sample sizes ranged from 1 to 16 individuals. Of those, 44 individuals were from Shi *et al*. (2013), the remaining 294 individuals were sampled in this study. We also included 12 individuals *M. przewalskii* (Zhang et al. 2020), and 10 individuals of *M. martensii* (Shi et al. 2013) in the phylogenetic analysis. These two species occur parapatrically with Mmon.

**Figure 1.**
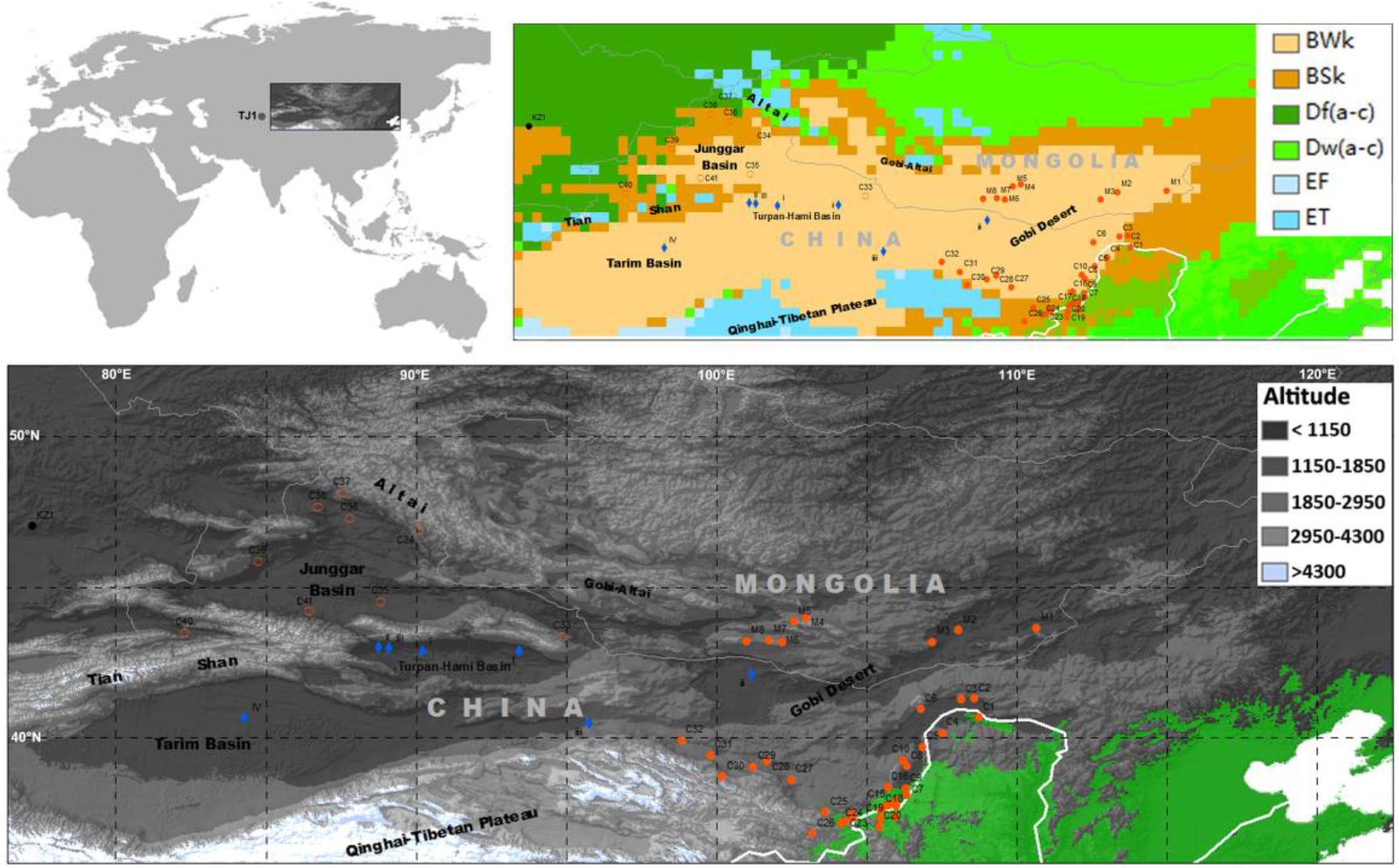
Geographic sampling of the desert scorpion *M. mongolicus* (Mmon). The sampled sites were plotted on a digital elevation map of the studies areas. The sampled sites were also projected on the map of Köppen-Geiger climatic classification (inserted, upright). BWk: arid, desert, cold; BSk: arid, steppe, cold; Df(a-c): cold, without dry season; Dw(a-c) cold, dry winter; EF: polar, frost; ET: polar, tundra. Orange dots, the Gobi population of Mmon; orange circles, the Junggar population of Mmon; blue diamonds, *M. prezwalskii*. Mmon and *M. prezwalskii* are typical arid species occurring widely in arid ecosystems (BWk and BSk). Green area in the low right represents suitable distribution range of *M. martensii*.

Genomic DNA was extracted from preserved tissues using a modified phenol-chloroform extraction procedure (Zhang and Hewitt 1998). All individuals were sequenced for the mitochondrial cytochrome *C* oxidase subunit I (mtCOI) gene and a subsample were sequenced for three nuclear loci, viz: nuclear protein kinase (PK), Serin-type endopeptidase (STE) and Serin proteinase inhibitor (Spn2) genes. The primers, PCR profiles and sequencing protocols followed Shi *et al*. (Shi et al. 2013). All specimens and DNA extracts were deposited at the Laboratory of Molecular Ecology and Evolution, Institute of Zoology (MEE-IOZ), Chinese Academy of Sciences, Beijing. The unique haplotypes have been deposited in GenBank under accession numbers xxxxxxxx-xxxxxxxx.

### Phylogenetic analyses and divergent time estimate

Sequences were aligned using MUSCLE (Edgar 2004). Phylogenetic analyses were performed using both maximum likelihood (ML) and Bayesian methods. The best-fit model of DNA evolution was selected using Modeltest 3.7 program (Posada and Crandall 1998) according to the Akaike information criterion (AIC). ML analysis was carried out using PHYML 3.0 (Guindon et al. 2010) with SPR tree search operation. Bootstrap with 1000 replicates was performed to assess topological robustness. Bayesian analysis was carried out with MrBayes 3.2 (Ronquist et al. 2012). Analyses were initiated with random starting trees and run for 10^6^ generations with four Markov chains employed. Trees were sampled every 100 generations and the ‘temperature’ parameter was set to 0.2. The first 30% trees were discarded as burn-in after checking for stationary and the convergence. Median-joining network (Bandelt et al. 1999) was constructed using the Network 4.6 .1.2 (http://www.fluxus-engineering.com/) to visualize intraspecific relationships among distinct haplotypes.

Divergence time was estimated with a relaxed molecular clock approach implemented in BEAST 1.10.4 (Suchard et al. 2018). We combined the most diverged mtCOI haplotypes revealed here with our previous alignment into a new data set with 53 sequences and estimated divergent times with the same calibrating method (Shi et al. 2013). The rate change was explicitly modeled using uncorrelated lognormal distribution across tree. Using a GTR +Γ model and a Yule process for speciation prior, we performed 100 million Markov chain Monte Carlo (MCMC) searches, sampling once every 1000 generations. Stationary and the convergence of the MCMC chains were checked with Tracer 1.7 (Rambaut et al. 2018). Maximum clade credibility (MCC) tree, posteriors, means and 95% highest posterior densities (HPDs) of ages of nodes were identified and annotated using TreeAnnotator 1.10 (Suchard et al. 2018).

### Phylogeographic diffusion analysis

Current and predicted suitable distribution area of Mmon stretches continuously across the entire studied region (Shi et al. 2015). There are no obvious potential barriers that can be enrolled to define discrete areas, and thus it is arbitrary to divide species range into discrete operational areas *a priori*. Therefore, we inferred spatial diffusion of genealogies using a continuous Bayesian phylogeographic method which models geographic spread of genealogies across spaces with continuous diffusion processes conditioned on sampling location defined as bivariate trait representing latitude and longitude for each accession (Lemey et al. 2010). This analysis was only based on the mtCOI data due to limited spatial sampling and low genetic variation of the nuclear genes. We first selected the best fit diffusion model by performing marginal likelihood estimation using generalized stepping-stone sampling on three available relaxed random walk models, as well as the time-homogeneous Brownian motion process. The Bayesian phylogeographic diffusion analysis was performed using BEAST v. 1.10.4 (Suchard et al. 2018). MCMC were run for 50 million generations with trees and parameters sampled every 5,000 generations. MCMC samples were summarized using TreeAnnotator. The results of phylogeographic reconstruction were visualized in SpreaD3 (Bielejec et al. 2016).

### Demographic inferences

Historical changes in the effective population size were assessed using two approaches. First, we used summary statistic tests, *D* (Tajima 1989), *Fs* (Fu 1997) and *R_2_* (Ramos-Onsins and Rozas 2002). Tajima’s *D* test is based on the number of segregating sites, and Fu’s *Fs* test uses information from the haplotype distribution, while Ramos-Onsins and Rozas’s *R_2_* test uses information of the mutation frequency. These statistics have high power to detect population expansion for non-recombining regions of the genome under a variety of different circumstances, either when sample size is large (*Fs*) or when sample sizes is small (*D*), and the number of segregating sites is low (*R_2_*). The significance of each test was assessed by generating null distribution from 10,000 coalescent simulations in DnaSP version 5.10.01 (Librado and Rozas 2009). Significantly large negative *D* and *Fs* values and significantly positive *R_2_* value were taken as evidence of population expansion. Second, demographic changes over time were explored by coalescent-based Bayesian skyline plot (BSP) using BEAST 1.10.4 (Suchard et al. 2018). Unlike above tests, which are based on summary statistics, BSP makes use of all the historical information contained within a sample of DNA sequences from a given population. The time estimated from molecular dating was used to calibrate BSP analyses. We ran the analysis for 5 × 10^8^ iterations, treating the first 10% iterations as burn-in, and then sampled genealogies and model parameters every 5, 000 generations for subsequent analysis.

### Ecological Niche Modeling

A total of 89 occurrence records for Mmon were used for ecological niche modeling (ENM) with MaxEnt version 3.3.3k (Phillips et al. 2006). We removed localities that were within distances of 30 km to each another to reduce the effect of sampling bias. We used the bioclimatic variables for both the present and past (21 kyr B.P., based on MIRCO models) at 2.5 arc-minute resolution (c. 5×5 km) in the WorldClim data set (Hijmans et al. 2005). The present bioclimatic layers were masked to region spanning from 35.8 to 55.4 °N and from 76.4 to 122.5 °E, to avoid sampling unrealistic background data and thus inflating the strength of predictions. The pairwise Pearson’s correlations between the climatic variables were used to restrict climatic variables. We removed highly correlated (|*r*| ≥ 0.80) climatic variables before ENM.

MaxEnt was run with a convergence threshold of 10^-5^ and maximum number of iterations of 10,000 with 25% of localities for model testing for 10 replicates with subsample. The model was projected to past conditions with clamping. Model performance was assessed via the area under the ROC (receiver operating characteristic) curve (AUC) statistic and the importance of variables was assessed by jackknife tests. AUCs > 0.75 are considered adequate for species distribution modeling applications (Fielding & Bell, 1997; Pearce & Ferrier, 2000). We employed two threshold rules, the maximum training sensitivity plus specificity threshold (MTSS), and the equate entropy of thresholded and original distributions logistic threshold (ENT), for converting continuous models to binary predictions. The MTSS is considered to most accurately predict presence/absence (Jiménez-Valverde and Lobo 2007). The ENT is known significantly over-predicts species ranges (Worth et al. 2014).

## Results

### Phylogenetic relationships and divergence times

The total 338 mtCOI sequences collapsed into 62 unique haplotypes, of which one haplotype was from Tajikistan (TJ01), one haplotype was from Kazakhstan (KZ01), and all other 60 haplotypes were from northwest China and south Mongolia. Combining with 12 samples of *M. przewalskii*, 10 samples of *M. martensii*, 6 representatives for another three species of *M. eupeus* complex and two outgroups, the final dataset includes 92 mtCOI haplotype sequences. The results of ML analysis and Bayesian inference were highly congruent. The ML phylogeny inferred from mtCOI dataset is shown in Figure 2A. All samples of Mmon from China and Mongolia clustered within an unresolved star clade which coalesced with TJ01 from Tajikistan with strong supports (95/1.00 for ML/Bayesian, respectively). We recognize this clade as Mmon. Within the Mmon clade, all haplotypes except one from the Junggar Basin (Xinjiang) formed a sub-lineage but was not strongly supported. This result was mirrored by the haplotype network analysis. Haplotypes from the Junggar Basin were closely related to those from the Gobi region but no haplotype was shared between two regions (Figure 2B). The Mmon clade coalesced with KZ01 and then *M. thersites* and *M. phillipsi*, forming a monophyletic branch (71/0.99) including exclusively members of the *M. eupeus* species complex. Monophyly for both *M. prezwalskii* and *M. martensii* was supported. These two species were more closely related each other than to any member of the *M. eupeus* complex.

**Figure 2.**
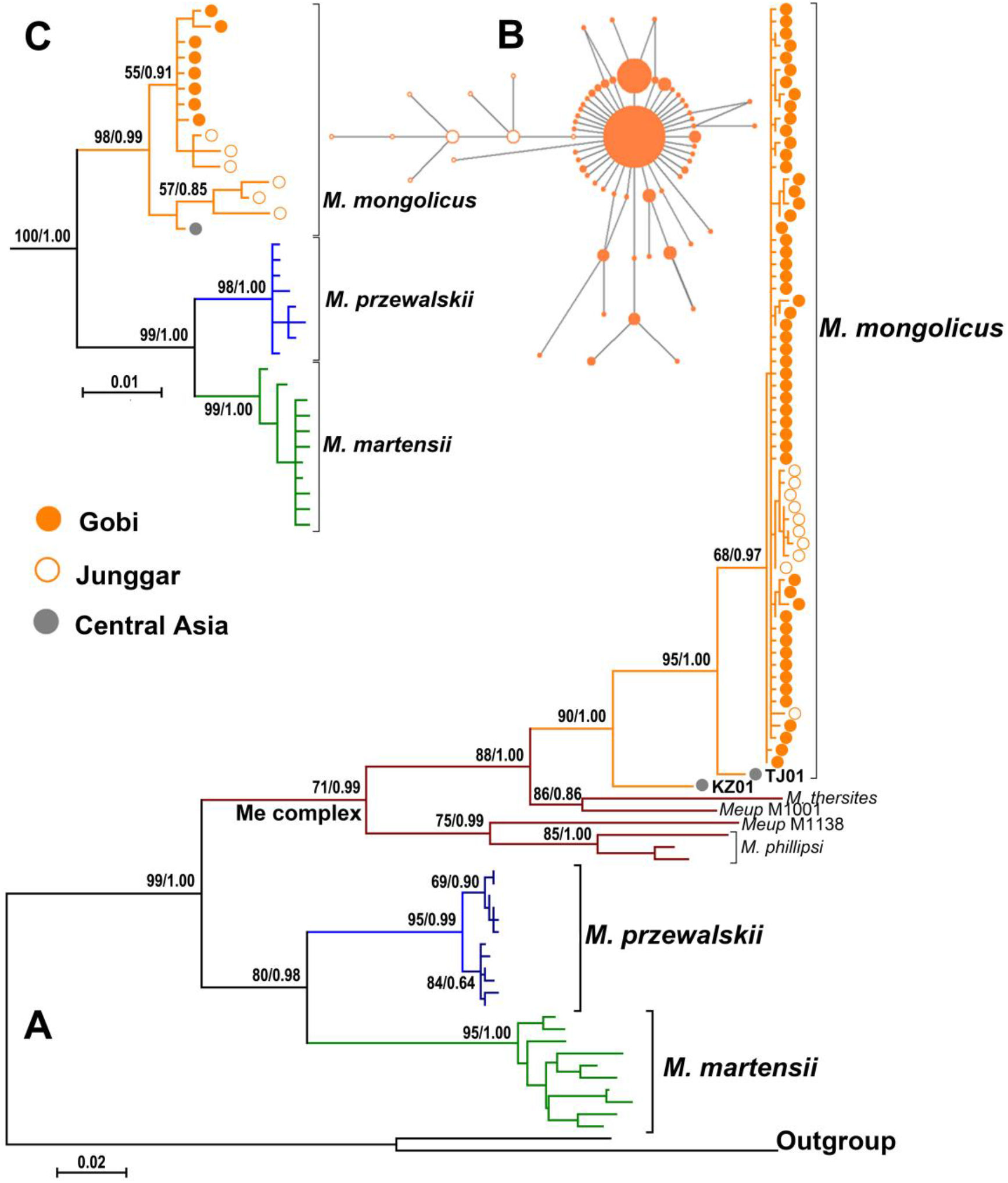
The desert scorpion *M. mongolicus* (Mmon) is the most derived lineage of the *M. eupeus* species complex. (**A**) Maximum likelihood tree inferred from the mitochondrial DNA data (mtCOI). (**B**) Median-joining network of Mmon mitochondrial haplotypes. Branch lengths are proportional to the number of substitutions occurred between haplotypes and circle sizes proportional to the number of individuals sharing the haplotypes. (**C**) Maximum likelihood tree inferred from three concatenated nuclear genes (PK, STE and Spn2). Numbers at the nodes represents the bootstrap support value and Bayesian posterior probabilities. Me complex, *M. eupeus* species complex.

The concatenated matrix for three nuclear loci (PK, STE and Spn2) included 33 samples. The results of phylogenetic analyses were congruent with the mitochondrial phylogeny as far as the interspecies relationship was concerned (Figure 2C). However, the nuclear phylogeny provided more resolved relationships within Mmon. Two moderately supported subclades were observed. One subclade includes three samples from the Junggar Basin and TJ01 from Tajikistan. The other subclade includes all samples from the Gobi region and three clustered samples from the Junggar Basin.

The results of divergence time estimation were summarized in Table 1. The time of the most recent common ancestor (*T_MRCA_*) of Mmon was estimated to be 1.36 Ma (95% HPD, 0.78–1.97 Ma) and the Junggar-Gobi lineage coalesced at 0.54 Ma (0.30-0.80 Ma). The divergence time between Mmon and KZ01 is 3.37 Ma (2.12-4.54 Ma), while that between *M. prezwalskii* and *M. martensii* is 6.27 Ma (4.35-8.37 Ma). A further split within *M. prezwalskii* occurred at 1.00 Ma (0.56-1.51 Ma). Divergence between the *M. eupeus* complex and the *M. prezwalskii-M. martensii* clade occurred around 10.56 Ma (8.33-12.70 Ma).

**Table 1.** Divergence time estimates generated by the relaxed molecular clock approach based on *mtCOI* data. Means and the corresponding 95% highest posterior density (HPD) intervals for times to most recent common ancestor (*T_MRCA_*) were given in million years for each lineage as shown in Fig. 2.

| MRCA | Mean | HPD lower | HPD upper |
| --- | --- | --- | --- |
| *M. mongolicus* (Mmon)-KZ01 | 3.27 | 2.12 | 4.54 |
| *M. mongolicus* (Mmon) | 1.36 | 0.78 | 1.97 |
| Junggar-Gobi lineage | 0.54 | 0.30 | 0.80 |
| *M. przewalskii* | 1.00 | 0.56 | 1.51 |
| *M. przewalskii*-*M. martensii* | 6.27 | 4.35 | 8.37 |
| Mmon- *M. martensii* | 10.56 | 8.33 | 12.70 |

### The spatiotemporal diffusion of Mmon

A Bayesian phylogeographic reconstruction of ancestral distribution and spatial diffusion of genealogies through time supports that Mmon was originated in Central Asia and dispersed eastward step by step, and finally to the Gobi regions. Three snapshots illustrating the major phylogeographic dispersal events are shown in Figure 3. The most likely ancestral distribution of the last most recent common ancestors of Mmon located in an area to the northwest of the Tianshan, somewhere near the boundary between Tajikistan and Kazakhstan. The ancestral population moved eastward and reached the Junggar Basin around 0.8 Ma, and then underwent regional dispersion within the Junggar Basin. Around 0.4 Ma, Mmon dispersed further east to the southern part of the Gobi, followed by another regional expansion in areas near the Tengger Desert and the Badan Jaran Desert, striding Gansu, Inner Mongolia and Ningxia. It was a very recent event (<0.2 Ma) that Mmon population expanded northward to the Gobi region of southern Mongolia. During this time, a northward expansion event was also occurred in the Junggar basin which gave rise to the Altay population. It appeared that regional population expansions have persisted until the present day, which gives rise to current trans-Gobi distribution for this species.

**Figure 3.**
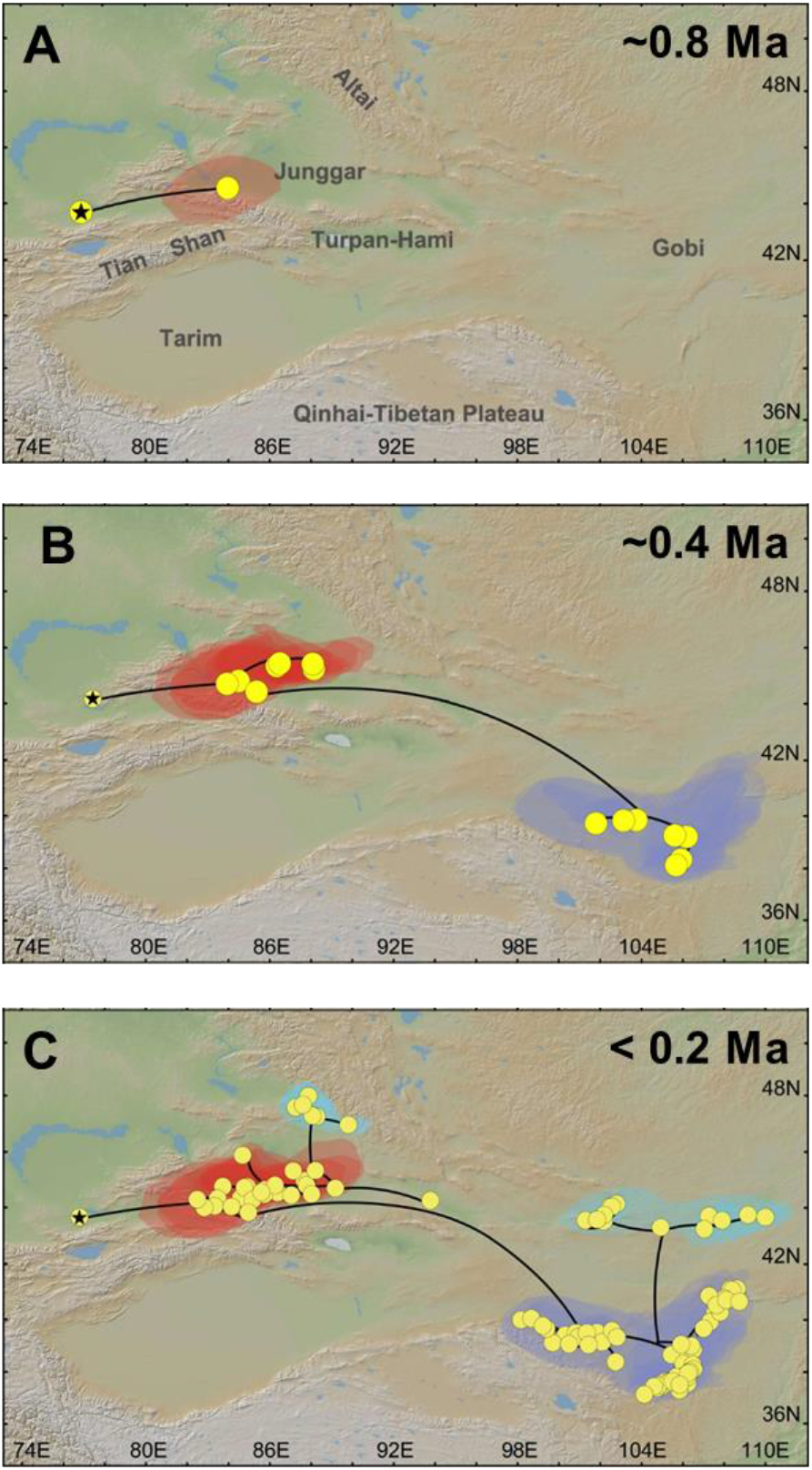
Spatiotemporal reconstruction of the geographic dispersal of *M. mongolicus* (Mmon) using the Bayesian phylogeographic method. Shown here are three snopshots shown that Mmon was originated in Central Asia (**A**), then dispersed eastward step by step (**B**), and finally to the Gobi regions (**C**). Geographic spread of genealogies across spaces was modelled with a continuous diffusion process governed by the relaxed random walk. Circles represent nodes of the maximum clade credibility phylogeny. Shaded areas represent 80% highest posterior density interval which depicted the uncertainty of phylogeographic estimates for the nodes.

### Demographic history

All the values for Fu’s *Fs* were negative and significant (*P* < 0.01), and all the values of Ramos-Onsins and Rozas’ *R_2_* were small (< 0.12), positive and significant (*P* < 0.05) for populations of Mmon (Table 2). The values of Tajima’s *D* for all except the Junggar population were negative and significant (*P* < 0.01). All these tests for *M. prezwalskii* were not significant (*P* > 0.3, data not shown). The variations of effective population size through time are shown by Bayesian skyline plots in Figure 4. For Mmon, the curve of BSP can be visually divided in to three phases. Before 35 ka, Mmon population were very stable. From 34 to 14 ka, Mmon population size increased sharply, even during the LGM around 21 ka. Since 14 ka until to present, the BSP curve of Mmon gradually plateaued off (Figure 4A). The population size of *M. prezwalskii* increased slightly since 400 kya and leveled off approaching the present day (Figure 4B).

**Figure 4.**
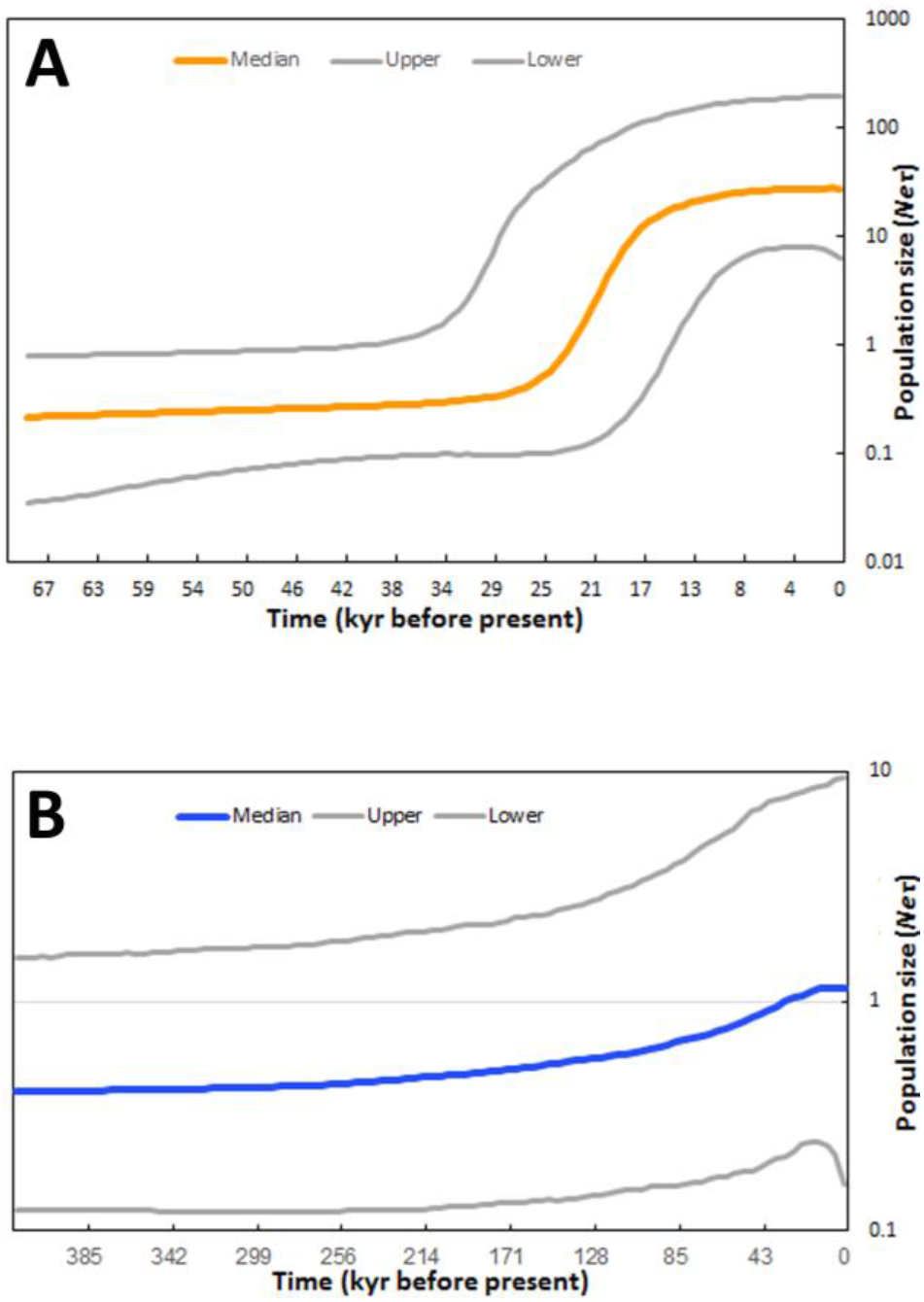
Bayesian skyline plots derived from samples of mtCOI sequences for *M. mongolicus* (**A**) and *M. prezwalskii* (**B**). The medians and their 95% highest posterior density intervals of effective population size (*N_e_τ*) were plotted against time. The *x* axis represents time in thousand years (kyr) before present. The *y* axis is the log scaled effective population size (*N_e_τ*). Population size for both species increased during the LGM about 21 kyr ago.

**Table 2.** Summary of genetic statistics of mtCOI data for *M. mongolicus* (Mmon). Here given are sample size (N), number of haplotypes (h), hapotype diversity (Hd), nucleotide diversity (π), Tajima’s *D*, Fu’s *Fs*, and Ramos-Osins & Rozas’ *R_2_* statistics. ** *P* < 0.01, ** *P* < 0.05

| Populatio | N | h | Hd | $\pi$ | *D* | *F<sub>s</sub>* | *R<sub>2</sub>* |
| --- | --- | --- | --- | --- | --- | --- | --- |
| **Mmon** | 336 | 60 | 0.7189 | 0.0023 | -2.3955** | -92.0339** | 0.0761** |
| **Junggar** | 19 | 7 | 0.7485 | 0.0019 | -1.0758 | -3.1589** | 0.0909* |
| **Gobi** | 317 | 53 | 0.6849 | 0.0020 | -2.4037** | -28.8463** | 0.0122** |
| South | 255 | 45 | 0.7399 | 0.0023 | -2.3120** | -59.5270** | 0.0798** |
| North | 62 | 10 | 0.3246 | 0.0006 | -2.1978** | -11.9593** | 0.1198** |

### Temporal changes of suitable distributional areas

After removing highly correlated (|*r*| ≥ 0.80) climatic variables, six climatic variables (BIO1, annual mean temperature; BIO7, temperature annual range; BIO9, mean temperature of driest quarter; BIO12, annual precipitation; BIO15, precipitation seasonality; BIO19, precipitation of coldest quarter) were used for final ecological modeling. These variables seem to be biologically meaningful for Mmon considering its ecology and the climatic regime of the studied regions (Shi et al. 2007). The distributional models generated with these six climatic variables maintained high AUC statistics (average 0.928 ± 0.023). The potential distribution of Mmon averaged over 10 models for the present climate are shown on Figure 5A. It encompassed the entire Junngar Basin and the Gobi region, with a narrow belt along the Hexi Corridor connecting these major areas. The northern rim of the Tarim Basin was also suitable for distribution of Mmon. However, both the Tianshan and Altai mountains were obviously unsuitable. Figure 5B shows the Palaeo-projection of ENMs at the LGM. It indicated sharp contraction of suitable range at this time, resulting in two major disjoined areas in the Junggar Basin and to the south of Gobi, respectively. In those regions, extensive southward retreat of potential niche occurred in the Gobi region, with the northern Mongolian Gobi becoming completely unsuitable and the Badian Jaran-Tengger sand deserts being the center of distribution. In the western part, the suitable area shrank towards the heart of the Junggar Basin. Of the 6 climate variables used, annual precipitation (BIO12) contributed to the MaxEnt models most (with a mean relative contribution of 47.2 %), flowed by annual mean temperature (BIO1, 31.1%), and mean temperature of driest quarter (BIO9, 16.4%). The three remaining variables collectively contributed 5.4% to the model.

**Figure 5.**
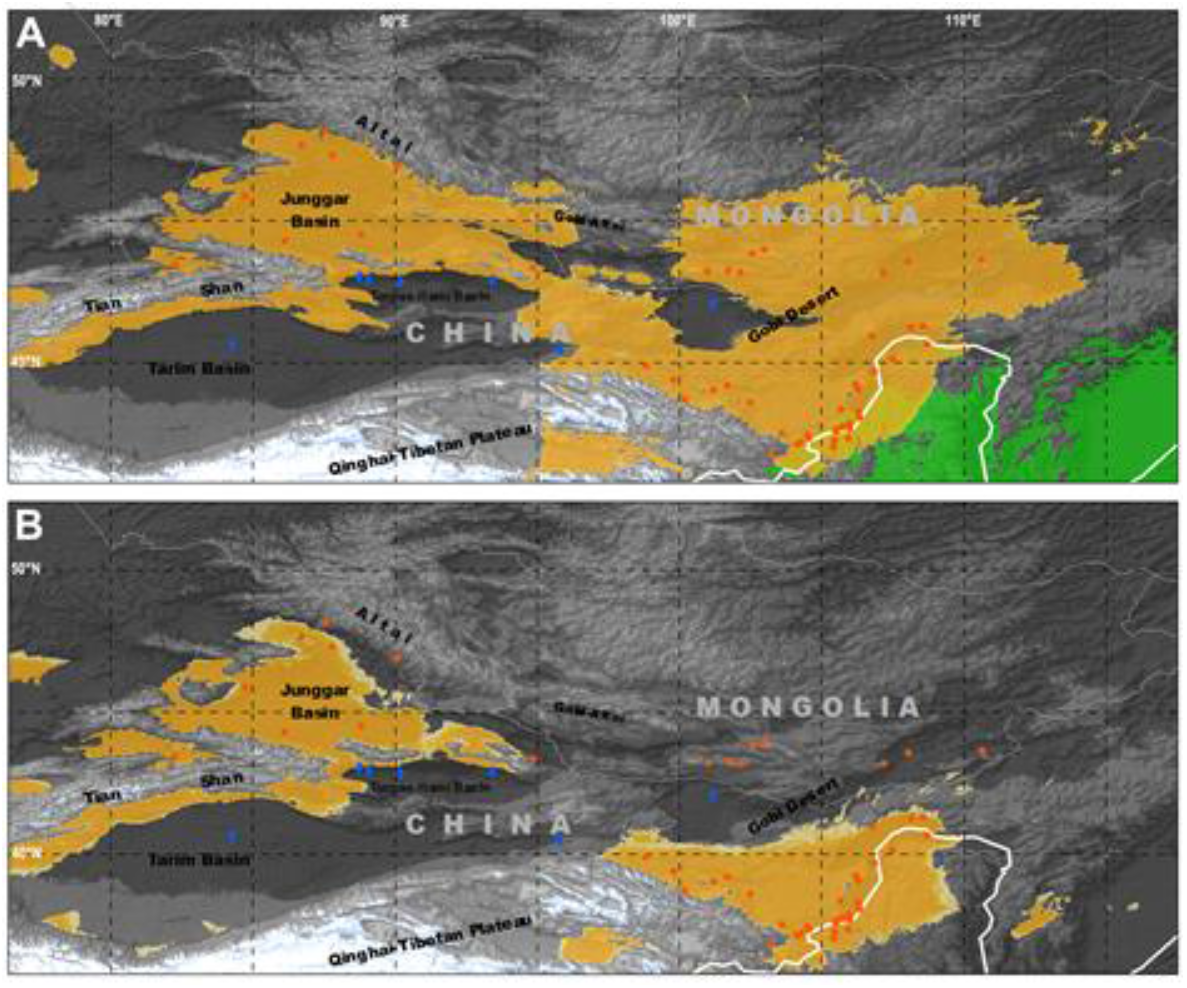
Potentially suitable distribution areas for *M. mongolicus* (Mmon) at present (**A**) and during the LGM (**B**). Ecological niche model generated under current conditions was hind projected to climatic condition at the LGM using Maxent. Orange areas represent the inferred range based on the MTSS threshold, and the yellow areas are the inferred range according to the ENT threshold.The green area in the low right represents suitable distribution range of *M. martensii*.

## Discussion

Past climatic oscillations have triggered recurrent reorganization of biota at all levels and scales, from genes to ecosystems (Davis and Shaw 2001; Hewitt 2000). In response to the climatic changes, populations must migrate to remain within their ecological niche or adapt to tolerate the challenges in order to survive and thrive (Davis and Shaw 2001; de Lafontaine et al. 2018; Fernandez et al. 2021). Otherwise, they will risk extinction. Scorpions of *Mesobuthus* (*sensu* Gantenbein et al. 2003) faithfully followed this general rule, thus emerging as an excellent model for studying biogeographical evolution in drylands of Asian interior (Graham et al. 2019; Shi et al. 2013). Since the late Miocene, this group has undergone extensive diversification and biogeographic evolution driven by regional tectonics (Mirshamsi et al. 2010; Mirshamsi et al. 2011) and climate changes (Shi et al. 2013), which gave rise to more than two dozen species widely spread in mid latitude Eurasia with Central Asia being center of species diversity (Fet et al. 2021; Fet et al. 2018; Fet et al. 2000; Graham et al. 2019). Currently, at least five species of *Mesobuthus* occur in China which was hypothesized as an outcome of multiple ‘Out of Central Asia’ evolution of the ancestral species (Shi et al. 2013). The Gobi population of Mmon was suggested to have undergone very recent out-of-Central Asia move in late Pleistocene (Shi et al. 2013). In the present study, we first define the route via which ancestral population of Mmon dispersed out of Central Asia and reached the Gobi, then tested the expansion-contraction model that might have incurred by climate-driven dispersal events, and finally explored the environmental context that has sustained such phylogeographic history.

### The Junggar Basin as the route out of Central Asia for the desert scorpion

Our study provides several lines of evidence that Mmon dispersed out of Central Asia and colonized the Gobi via the Junggar Basin along the northern side of the Tianshan mountains. Previously, we have shown that the ancestral distribution area of the *M. eupeus* complex was most likely in Anatolia and the ancestral population of Mmon arose somewhere in Central Asia (Shi et al. 2013). In the context of regional tectonic history and current topographic configuration such as the high mountain ranges (*i.e*. Tianshan and Pamir), it was possible for ancestral Mmon to disperse to the Gobi either through the Junggar Basin to the north of Tianshan (Junggar route) or through the Tarim Basin to the south of Tianshan (Tarim route), or by taking both routes simultaneously. Our results suggest that the Junggar route was the most likely one.

With extensive samples from China and Mongolia, we identified a single geographically coherent Mmon lineage which coalesced around 0.54 Ma and diverged from its most closely related lineage in Central Asia (TJ01) around 1.36 Ma (Table 1). Given ancient uplift of the Tianshan mountains forming a formidable geographical barrier (Figure 5), we would expect a much deeper genetic divergence than we observed here between the Central Asian population (*e.g*. TJ01) and the Gobi populations, if the desert scorpion dispersed out of the Central Asia along the Tarim route. During the late Miocene, most part of the Tianshan mountains have reached a critical height to be an effective geographical barrier (Yin 2010). The single lowland gateway connecting Central Asia and the Tarim Basin closed at 5.3 Ma due to the collision of the Tianshan orogen and the Pamir salient (Sun et al. 2015). Thus, if Mmon had dispersed via the Tarim route, it would imply a divergence date of around 5.3 Ma between the Central Asia population and the Gobi population. This date is more than three times earlier than the date we have estimated here (1.36 Ma, Table 1). Thus, the Tarim route for Mmon dispersal out of Central Asia can be largely excluded.

Indeed, Bayesian phylogeographic reconstruction clearly supported the Junggar route out of Central Asia for Mmon (Figure 3). During the past 1.36 million years, Mmon has been on an active eastward move. The ancestral population entered the Junggar Basin around 0.8 Ma and reached the Gobi region round 0.4 Ma. We noticed that such a colonization route from Central Asia was not limited to the scorpion species we have studied. Similar phylogeographic patterns have been reported in the agamid lizards (*Phrynocephalus* sp.) (Melville et al. 2009), and the rapid racerunner (*Eremias velox*) (Liu et al. 2019), although the times of colonization might differ. These evidences suggest that the Junggar Basin have constituted an important corridor for fauna exchange between Central Asia and the Gobi regions.

### Demographic expansion of the desert scorpion decoupled from range expansion during the LGM

A typical response of temperate species to past climate changes is demographic fluctuations. The star-shaped genealogy (Figure 2AC), the excess of rare mutations and deficit of intermediate frequency mutation (Fig 2B), and the significant negative Tajima’s *D* and Fu’s *Fs* as well as significant small *R_2_* (Table 3), all supported population expansion in Mmon. The curve of BSP indicated that Mmon population underwent exponential growth sine about 35 ka and throughout the LGM (Figure 3A). Such a glacial maximum expansion phenomenon is in sharp contrast to the paradigmatic interglacial-expansion glacial-contraction model corroborated for temperate flora and fauna of North America and Europe (Hewitt 2004; Schmitt 2007). It is also inconsistent with the scenario for the mesic biota in arid regions of the other parts of the world, in which population contraction occurred during the LGM with the effect of aridification mirrored that of glaciation (Fujita et al. 2010). However, such a demographic pattern appears to be characteristic of arid-adapted species in northwest China. For example, population expansions during the LGM have been detected in the desert-dwelling gerbil *Meriones meridianus* (Wang et al. 2013), the desert plant species of *Lagochilus* (Meng and Zhang 2013) and *Zygophyllum xanthoxylon* (Shi and Zhang 2015). Glacial demographic expansions also occurred in arid-adapted species from other regions of the world, for instance, the grey butcherbird *Cracticus torquatus* (Kearns et al. 2014) from arid woodland of southern Australia, the lizard *Liolaemus darwinii* in the Monte Desert of South America (Camargo et al. 2013), and three sympatric beetle species (*Iliotona beyeri, Carcinops gilensis* and *Belonuchus* sp.) from the Sonoran Desert of North America (Pfeiler et al. 2013). Our results, along with these cases, substantiate that organisms’ ecological adaptations were an important determinant to their demographic response to past climate changes.

It has been repeatedly demonstrated in a plethora of animals and plants that demographic expansion was often accompanied by range expansion. However, Mmon appeared to be an exception. We observed that demographic expansion of Mmon was accompanied by range contraction. While Mmon experienced population expansion, ENM indicated that its geographical distribution was drastically reduced during the LGM. That is, the suitable climate niche shrank toward the southern Junggar Basin and the southern Gobi (Figure 5). Nevertheless, such a scenario was reasonable when considering collectively the regional climate context and the scorpion ecology. Firstly, climate change in these regions seemed moderate at the LGM. Only limited glacial expansion occurred in Tianshan and other nearby high mountains during the LGM (Lifton et al. 2014). Secondly, Mmon is an arid-adapted species, and thus aridification associated with glaciation would exert limited stress to its persistence. On the contrary, the consequential expansion of desert environments might have opened up an array of new habitats that became accessible to Mmon. Besides, the extent of sand dune areas was expanded during glacial periods (Dong et al. 2013; Lu et al. 2013), which would provide Mmon suitable matrix for burrowing, by which scorpions could effectively buffer the climate extremes. As a consequence, although the overall size of their niche area was reduced during the LGM, the localized effective niche spaces might have expanded with the development of the desert ecosystems. Third, given that Mmon is the only scorpion species that has colonized the Gobi region rather recently (Shi et al. 2013) and that scorpions can maintain extremely high biomass in arid communities (Polis and Yamashita 1991), it could be expected that the population density would be far below the carrying capacity of the habitats even at the LGM, allowing proliferation of Mmon populations. In summary, the unique ecological adaptation of the desert scorpion and the distinct regional climate regime should have collectively molded the observed demographic pattern of Mmon in Asian drylands.

### The environmental context for phylogeographic evolution in Asian drylands

The evolutionary history of the desert scorpion Mmon reconciled well with the history of environmental changes in Asia drylands. The single geographically coherent lineage from China and Mongolia diverged from its most closely related lineage in Tajikistan around 1.36 Ma (0.78-1.97 Ma). Around this time, Central Asia become increasingly arid (Ding et al. 2002; Yang and Ding 2006) and northern China experienced an abrupt transition to sharply intensified winter monsoon (Ding et al. 1994; Liu et al. 1999), accompanied by expansion of desert environment (Ding et al. 2005). Study on the loess in Tajikistan also showed an increased aridity in Central Asia since 0.85 Ma (Ding et al. 2002; Yang and Ding 2006). Eolian loess deposited on the northern slope of the Tianshan indicated that the extremely dry climatic conditions was present in the Junggar Basin at least by 0.8 Ma and that the desert there experienced two dramatic periods of expansions at ~0.65 Ma and 0.5 Ma, respectively (Fang et al. 2002). These environmental changes might have provided the desert scorpion Mmon an opportunity to expand further eastward out of the Junggar Basin. To more eastern region, Badain Jaran and Tengger sand deserts were in place around 0.85 Ma (Guan et al. 2011; Li et al. 2014). The arid desert landscape occurred on the alluvial plain of the Gobi-Altai mountains no later than 0. 42 Ma (Lü et al. 2010). The sudden radiation of Mmon and its out of Central Asia dispersal (around 0.54 Ma, 0.3-0.8 Ma) was in the frame of these aridification events. These stepwise expansion of desert environment (Ding et al. 2005) should have opened up suitable habitats for Mmon to disperse eastward and finally reached the Gobi region. As revealed by hindcast ENM, large suitable areas existed both in the Junggar Basin and in the Gobi region although become disjunct at LGM (Figure 5B). Such large refugial regions would have sustained demographic expansion even during the LGM (Fig 4).

In summary, by revealing a specific phylogeographic history of the desert scorpion *M. mongolicus*, which was relevant to its biology and reconciled to regional climate change regime, we have demonstrated that (1) the eastward dispersal of the desert scorpion out of Central Asia to the Gobi was driven by intensified aridification events, and most likely through the Junggar Basin which have constituted an important corridor for fauna exchange between Central Asia and the Gobi regions, (2) the demographic expansion of the desert scorpion was accompanied by range contraction during the LGM, and thus (3) an organism’s ecological adaptation were an important determinant to their phylogeographic and demographic response to past climate changes. Future work directing to understanding the crucial limiting factors of species range from different geography will provide major scaffolds for integrating phylogeographic knowledge across spaces and times.

## Acknowledgements

We are grateful to the Plant Protection Research Institute, Mongolia, for logistic supports on field surveys. We thank Peng He, Hong-Bin Liang, Ke-Qing Song, Zhiyuan Su, Duo-Hong Wang, Zhi-Liang Wang for collecting scorpions.

## Author contributions

D-X.Z. and C-M.S. conceived and designed the project; C-M.S., X-S.Z., L.L and Y-J.J. collected the molecular data; C-M.S. performed the field work and analyzed data; C-M.S. and D-X.Z. wrote the manuscript.

